# Non-REM sleep facilitates consolidation of contextual fear memory through temporal coding among hippocampal neurons

**DOI:** 10.1101/2020.05.29.123356

**Authors:** Quinton M. Skilling, Brittany C. Clawson, Bolaji Eniwaye, James Shaver, Nicolette Ognjanovski, Sara J. Aton, Michal Zochowski

**Affiliations:** Biophysics Program, University of Michigan, 930 N University Ave., Ann Arbor, MI 48109; Department of Molecular, Cellular, and Developmental Biology, University of Michigan, 830 N University Ave., Ann Arbor, MI 48109; Applied Physics Program, University of Michigan, 450 Church St, Ann Arbor, MI 48109; Department of Physics, University of Michigan, 450 Church St, Ann Arbor, MI 48109

**Keywords:** memory consolidation, phase coding, oscillatory dynamics, sleep

## Abstract

Sleep plays a critical role in memory consolidation, although the exact mechanisms mediating this process are unknown. Combining computational and *in vivo* experimental approaches, we test the hypothesis that reduced cholinergic input to the hippocampus during non-rapid eye movement (NREM) sleep generates stable spike timing relationships between neurons. We find that the order of firing among neurons during a period of NREM sleep reflects their relative firing rates during prior wake, and changes as a function of prior learning. We show that learning-dependent pattern formation (e.g. “replay”) in the hippocampus during NREM, together with spike timing dependent plasticity (STDP), restructures network activity in a manner similar to that observed in brain circuits across periods of sleep. This suggests that sleep actively promotes memory consolidation by switching the network from rate-based to firing phase-based information encoding.

## Introduction

How the brain binds various sensory features of events into a neural representation for memory storage is a long-standing question in neuroscience. Available data suggest that initial memory encoding is driven by increases in activity (rate coding) among a specific population of neurons, often referred to as “engram neurons” [1-4]. Over time, these mnemonic representations are incorporated into more widely distributed networks in a process referred to as systems memory consolidation [5]. Recent data suggest that engrams are initially formed from heterogeneous neuronal populations with log-normal distributions of firing rates ranging over several orders of magnitude [6, 7]. This heterogeneity may reflect populations encoding different aspects of experience [7-9], which likely exhibit different dynamics during memory consolidation. A critical unanswered question is how these heterogeneous populations, distributed across the brain over vast synaptic distances, cooperate in the process of long-term memory storage.

Rate coding alone has limitations for long-term information storage *in vivo*. First, individual neurons have widely divergent baseline firing rates. This firing rate heterogeneity constitutes background “noise”, which could easily obscure firing rate changes in sparse populations of “signal” encoding neurons. Second, because individual neurons have a limited dynamic range over which their firing rates vary, rate coding has a limited capacity for integration over time. In other words, neurons’ peak firing rates would represent a ceiling, after which no additional information could be carried based on rate-coding alone. Third, changes in firing rate will alter spike timing-dependent plasticity (STDP) rules governing synaptic strengthening or weakening [10]. As neurons increase their firing rates, STDP will be either reduced overall (due to differences in firing rates between pre- and postsynaptic neurons), or biased toward potentiation [11]. Thus, the information-carrying capacity of STDP for the network is limited by firing rate increases. When these factors are taken together, it is clear that non-rate-based coding mechanisms must be invoked for long-term information storage (i.e., memory consolidation) in a heterogeneous neuronal network.

Oscillatory patterning of neuronal firing during sleep has been implicated in promoting synaptic plasticity and memory storage [12-19]. Network oscillations present in brain circuits during sleep have been implicated in promoting STDP by precisely timing the firing between pairs of neurons [14, 15]. Hypothetically, the transition to oscillatory dynamics between wake and sleep could constitute a switch between rate coding (useful during learning) and phase coding (which may promote consolidation). For the reasons outlined above, this switch from rate to phase coding (i.e. when neurons’ respective phase of firing on each cycle conveys the information about the network patterning) is essential for long-term information storage in brain networks.

Here, we combine computational modeling of a highly reduced CA1 hippocampal network with analyses of *in vivo* recordings from CA1 to investigate how hippocampal network structure and dynamics are affected during contextual fear memory (CFM) consolidation following single-trial contextual fear conditioning (CFC) [20]. CFM consolidation relies on *ad lib* sleep in the hours immediately following CFC [17, 21] – a process associated with enhanced delta- (0.5-4 Hz), theta- (4-12 Hz), and ripple-frequency (150-200 Hz) hippocampal activity in the hours following learning [22]. Recent *in vivo* work has shown CFM consolidation is disrupted when these oscillations are suppressed during post-CFC sleep deprivation [16, 17, 22]. Conversely, CFM can be rescued from disruption caused by experimental sleep deprivation when oscillations are driven optogenetically (via rhythmic activation of fast-spiking interneurons) in CA1 [17].

We show that regulation of resonance properties in the CA1 network model, implemented via changes in acetylcholine (ACh) signaling analogous to those occurring in NREM sleep, recreates a number of experimentally-observed phenomena. Within the model network, memory consolidation is state-dependent (occurring in a state analogous to NREM sleep), associated with augmented post-learning network oscillations and stabilization of functional communication between neurons, and blocked by suppression of inhibitory interneuron activity. Memory storage is associated with a switch between rate coding (during a wake-like state) and phase coding (during a NREM-like state) in the network. NREM phase coding drives STDP between network neurons which causes dramatic, differential changes in the strength of reciprocal connections between highly active vs. sparsely firing neuronal populations. These changes lead to differential changes in firing rate in the two populations – a finding consistent with experimental data from the CA1.

Together, our results show that successful memory consolidation requires the brain to switch between information coding schemes – a transition which occurs naturally through the sleep-wake cycle. Through this switch, functional network structures associated with engrams become more stable and robust for long-term information storage.

## Results

### Introducing a memory to the hippocampal network augments oscillatory dynamics and stabilizes functional connectivity patterns

We used hippocampal area CA1 following single-trial contextual fear conditioning (CFC) [*Na*, 24] as a model system to investigate how network changes during memory encoding affects subsequent network dynamics. We analyzed both *in vivo* recordings and *in silico* model network recordings of CA1 neurons to determine how functional network dynamics were affected by *de novo* memory formation [20, 22].

To investigate mechanisms involved in sleep-dependent aspects of memory consolidation, we simulated a reduced CA1 network model composed of excitatory principal neurons and inhibitory interneurons. For cells in the model, we used a conductance-based formalism (see **Methods**) incorporating a slow-varying potassium current which acts as a control parameter for neural spiking dynamics [25]. In the brain, this conductance is regulated by muscarinic ACh receptors [26]. During wake, high ACh blocks this current in excitatory neurons, which increases firing frequency responses to excitatory input and decreases firing synchrony with rhythmic input (i.e., type 1 excitability). Thus, during wake, neurons behave as integrators to the incoming stimuli, responding to changes in input by modulating their firing frequency. Low ACh during NREM sleep allows slow potassium current to play a larger role in membrane excitability, leading to spike frequency adaptation, reduced spike frequency response gain, and increased synchronization capacity (i.e., type 2 excitability) [25]. In this case, the neurons tend to spike with similar frequency across wider range of inputs, and lock to intrinsic or extrinsic oscillatory drive, especially if the oscillatory drive occurs at a resonant frequency.

In our model, sleep/wake dynamics are mimicked by switching excitatory neurons from type 2 membrane excitability (NREM sleep) to type 1 excitability (wake), resulting in neuronal excitability change from integrator type response during wake, into resonator type response during NREM sleep. Since the muscarinic response of interneurons to ACh is complex and heterogeneous [27, 28], we set inhibitory interneurons in the model to exhibit consistent type 1 dynamics (although permitting type 2 dynamics in interneurons yielded similar results).

Within the model we divided information storage into two phases. In the first, activation of engram neurons by external input results in strengthening of connections between them — a process we refer to as “learning” hereafter. In the second phase, off-line network reorganization is driven by STDP in a NREM-like state — which we refer to as “memory consolidation”. Within this framework, we first investigated how strengthening excitatory synaptic connections between a limited subset of neurons, an event analogous to initial learning, affects network activity patterns during subsequent sleep.

Comparing raster plots (**Figure 1A-B top)** and simulated local field potentials (LFPs; **Figure 1A-B bottom**) for the network when excitatory neurons exhibit type 2 (NREM) dynamics, before vs. after introduction of learning, reveals the emergence of well-defined oscillations (which are evident in periodic firing patterns of both excitatory and inhibitory neurons in the network). Learning also lead to an increase in network-wide low frequency spectral power (**Figure 1C**), consistent with previous *in vivo* work [17, 22] (effects of CFC on CA1 LFP activity during NREM sleep are shown in **Figure 1D)**. We hypothesized that this enhancement of coherent network oscillations (which has been widely reported in both human subjects and animal models following learning [15]) could drive STDP-based information storage in the network.

**Figure 1:**
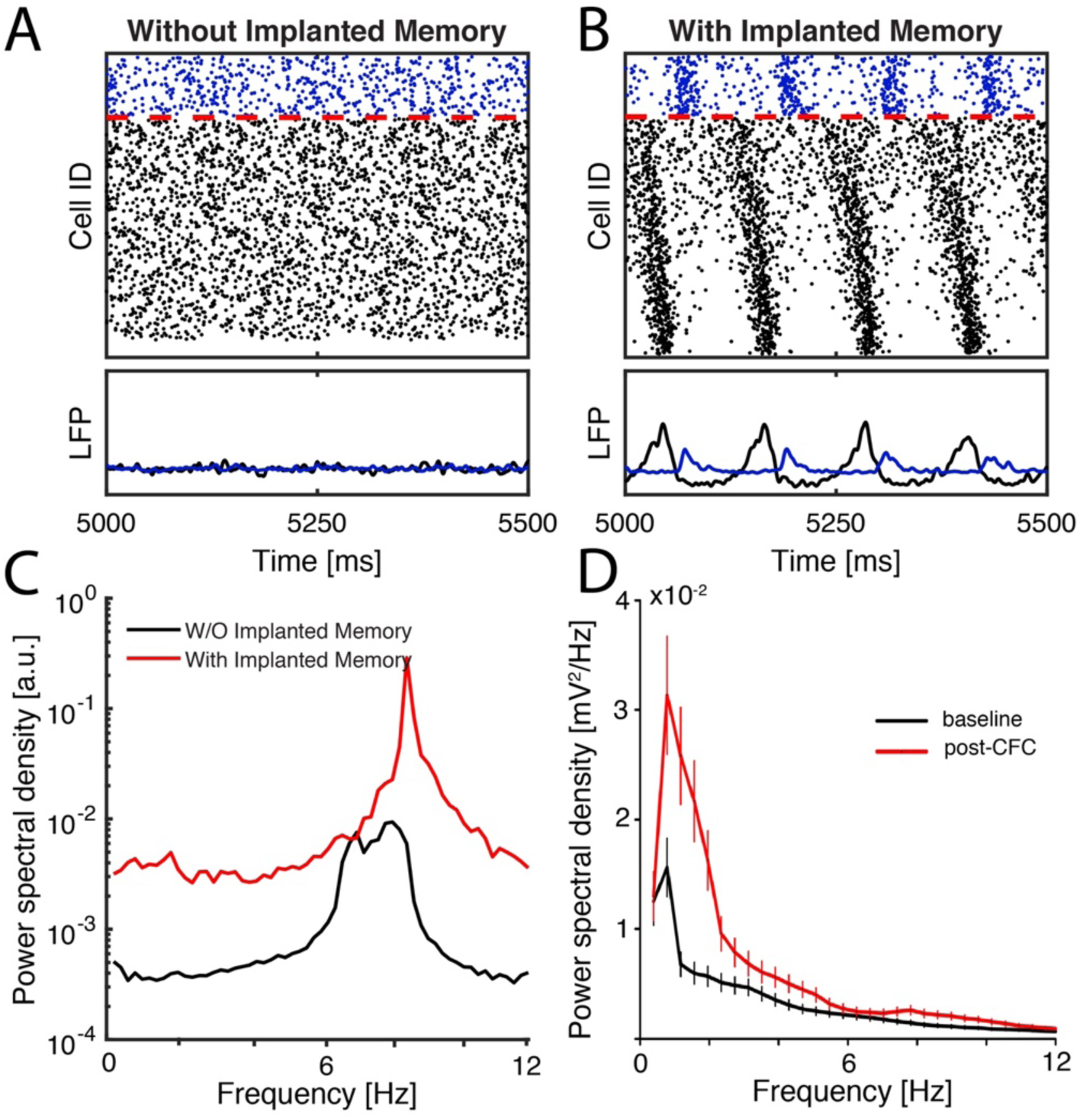
Model networks respond to sparse strengthening of excitatiory synapses through emergance of low-frequency rhythms and phase-locking. **A-B) SIMULATED DATA:** Raster plots (top) and cumulative signal (bottom) generated by inibitory (blue), and excitatory (black) neurons in a NREM-like (low-Ach) state before (**A**) and after (**B**) learning (i.e. 20 fold strengthening of synapses between pairs of excitatory engram neurons). Rasters in black and blue (below and above the red, dashed line) are excitatory and inhibitory neurons, respectively. **C)** Fourier transform of the excitatory network signal reveals an increase in spectral power at low frequencies; black line marks baseline; red line – spectral power after synaptic strenghtening. **D) EXPERIMENTAL DATA:** Fourier transform of CA1 LFP recordings during NREM sleep; spectral power is averaged over the first 6 h of baseline recording (black) and the first 6 h post-CFC (red).

### NREM hippocampal network stabilization *in vivo* predicts successful memory consolidation

To investigate how the emergent oscillations affects neurons’ firing relationships, we calculated the stability of functional connectivity between neurons, using the functional network stability metric (FuNS; see **Methods**) [29]. First, *in silico*, we simulated networks with differential exposure to learning (i.e. strengthening reciprocal excitatory connections among engram neurons as described above) and varying between type 1 and type 2 (wake and NREM sleep-like) network states. To model effects of post-learning sleep deprivation (SD), excitatory neurons were set to type 1 excitability in the presence of an implanted memory. To model interactions between learning and subsequent sleep, NREM was mimicked by setting excitatory neurons to type 2 excitability, in the presence or absence of an engram (Learning and Sham, respectively). When the engram was present in networks with type 2 excitability, the network exhibited more stable network dynamics over time (**Figure 2A**). Conversely, type 2 sham networks without an engram and type 1 networks with an engram showed no change in stability, even in the context of network STDP.

**Figure 2:**
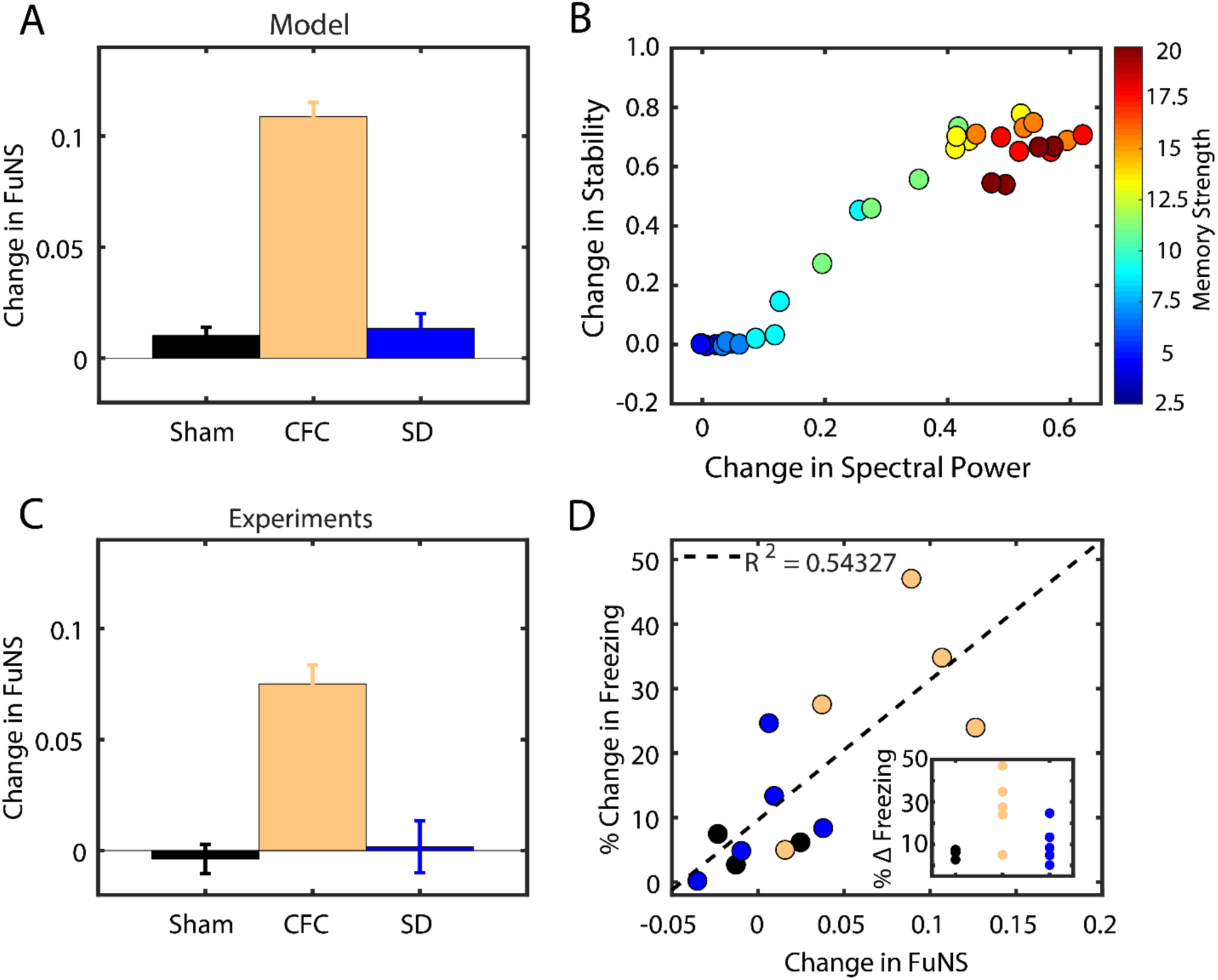
Functional network stability predicts future level of memory consolidation. **A) SIMULATED DATA:** Model predictions for the change in FuNS in each simulation group: Sham (NREM states without learning; n = 5) and SD (Wake states with learning; n = 5) show only marginal changes in FuNS whereas CFC (NREM states with learning; n = 5) show a maximal increase in FuNS. All error bars represent the standard error of the mean. **B) SIMULATED DATA:** Change in FuNS as a function of the change in spectral power density for 4-12 Hz band, for increasing the strength of those selected synaptic connections reveals a linear releationship. The color bar represents the magnitude of synaptic strength change from baseline. **C) EXPERIMENTAL DATA:** FuNS analysis of CA1 recordings following CFC or Sham conditioning (Sham conditioned — black [n = 3], CFC + SD — blue [n = 5], CFC + sleep – yellow [n = 5]). *Ad lib* sleep post conditioning leads to the greatest increase in FuNS. **D) EXPERIMENTAL DATA:** Change in FuNS for each mouse after CFC predicted its memory performance 24 h later (% freezing; raw values shown as **inset**). Line indicates best fit to data with R^2^ = 0.54.

To investigate how the magnitude of synaptic strength changes during initial learning subsequently affects FuNS and network oscillations during sleep, we measured these two quantities as a function of strength of connections between engram neurons. With increasing memory strength, we observed a positive linear relationship for increases in both FuNS and oscillatory power in the model network (**Figure 2B**). This suggests that learning in the network directly augmented both network oscillations and stabilization of functional connectivity patterns in subsequent NREM sleep.

We next tested how these features are affected during sleep-dependent consolidation of CFM *in vivo*. Mice either underwent single-trial CFC (placement into a novel environmental context, followed 2.5 min later by a 2-s, 0.75 mA foot shock; n = 5 mice), sham conditioning (placement in a novel context without foot shock; Sham; n = 3 mice), or CFC followed by 6 h of sleep deprivation (SD; a manipulation known to disrupt fear memory consolidation [17, 21, 30]; n = 5 mice). We measured changes in FuNS in these recordings after each manipulation by quantifying FuNS on a minute-by-minute basis over the entire pre- and post-training 24-h intervals and calculating their respective difference within each animal. Consistent with previous findings [20], we observed a significant increase in FuNS over the 24 h following CFC during NREM sleep (**Figure 2C**). In contrast, no change in NREM FuNS was seen in Sham mice or following CFC (during recovery NREM sleep, which is insufficient for CFM consolidation) in SD mice.

Group differences in NREM FuNS were reflected in the behavior of the mice 24 h post-training, when context-specific fear memory was assessed (inset **Figure 2D, inset**). Mice allowed *ad lib* sleep following CFC showed significantly greater freezing behavior when returned to the conditioned context than did Sham or SD mice. Moreover, CFC-induced changes in NREM-specific FuNS for individual mice predicted context-specific freezing during memory assessment 24 h later (**Figure 2D**). Thus, successful consolidation of a behaviorally-accessible memory trace *in vivo* is accompanied by increased NREM FuNS in the CA1 network.

Together, these results led us to hypothesize that oscillatory patterning could promote successful STDP-based consolidation of a hippocampal memory trace. We focus on this phenomenon in the following sections.

### Temporal organization of firing in network oscillations is a predictor of sleep-dependent firing rate reorganization

A series of recent studies have demonstrated that neuronal firing rates are renormalized across a period of sleep, with highly active neurons in a circuit reducing their firing rates, and sparsely firing neurons increasing their firing rates [6, 7, 31]. Sleep is essential for these firing rate changes, which do not occur across a period of experimental sleep deprivation [7]. We hypothesized that this phenomenon results from STDP driven by neurons phase-locking their firing to NREM sleep oscillations. Specifically, we predict that neurons that are highly active during wake will fire at an earlier phase within an oscillation than neurons with sparser firing. To test this, we first calculated phase of firing of every neuron with respect to LFP oscillations *in silico* (see **Figure 3A** and **Methods**). **Figure 3B** illustrates the relationship between model neurons’ phase of firing calculated during type 2 dynamics before learning as a function of the normalized frequency during type 1 dynamics. We observed that the fastest firing neurons during waking (type 1) fire earlier in the phase of the excitatory network oscillation during NREM sleep (type 2). This suggests that neurons take on a phase-based, temporal coding strategy during NREM network oscillations which reflects differences in firing rate present during wake.

**Figure 3:**
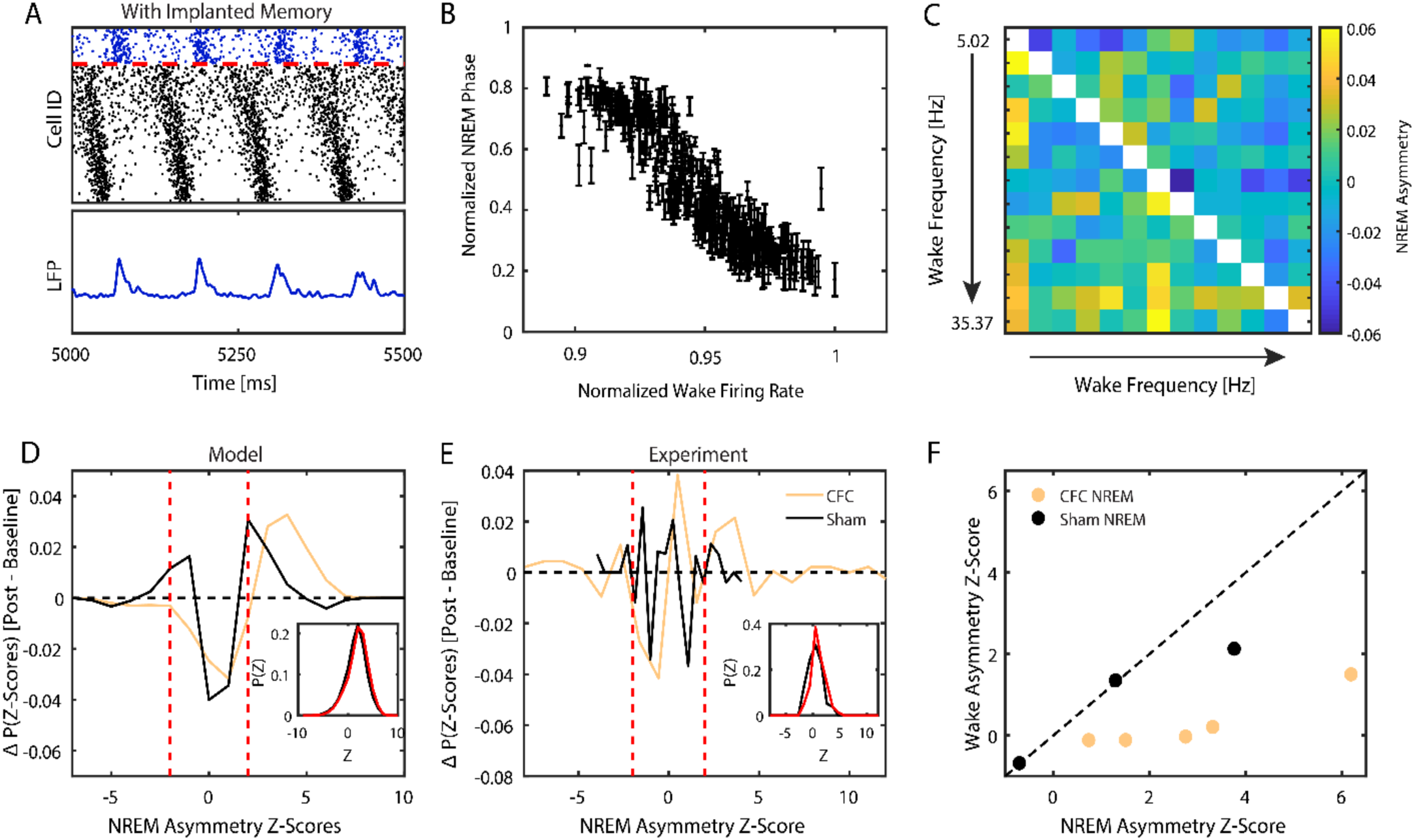
Mapping relative spiking frequency during wake onto phase relationships during NREM sleep. **A) SIMULATED DATA:** Calculation of mean phase of spiking relative to oscillatory signal exhibited by inhibitory population; peaks in the inhibitory signal are used as the starting points of the phase calculation. **B) SIMULATED DATA:** Normalized phase of firing in NREM versus normalized wake frequency of excitatory neurons reveals that the neurons firing with the highest frequency align with an earlier phase of the excitatory network population whereas slower firing neurons align with later phases. ***C-F)*** Analysis of NREM spiking asymmetry reveals enhanced wake frequency dependent temporal relationships between firing neurons after CFC. **C) SIMULATED DATA AND EXPERIMENTS:** Formation of pairwise spiking asymmetry matrix, *A*. Example: recorded activity from representative CFC mouse during NREM; rows and columns have been arranged by wake frequency; color denotes pairwise spiking asymmetry as defined in the methods section. **D) SIMULATED DATA:** Weighted average of differences in distributions (see inset for a representative example scenario; black—baseline, red—post) of pairwise Z-scores for models with type 2 dynamics and learning(yellow) and for models with type 2 dynamics and no learning (black). **E) EXPERIMENTAL DATA:** Weighted average of differences in distributions (see inset for example of representative mouse; black—baseline, red—post) of pairwise Z-scores for CFC mice (yellow) and Sham mice (black). **F) EXPERIMENTAL DATA:** significance (Z-score) of global NREM network asymmetry vs that of global wake asymmetry calculated for individual CFC (yellow) and sham (black) mice. Strong asymmetry is observed in NREM sleep but not in wake. Red dashed lines in (**D, E**) represent the significance cutoff of |*Z*| ≥ 2.

To investigate whether this phase-locking phenomenon can be observed experimentally, we compared *in silico* and *in vivo* network dynamics as a function of learning and sleep. We developed a metric to measure frequency-dependent phase-of-firing relationships between pairs of CA1 neurons by quantifying their spike timing asymmetry within bursts of activity (for a full description, see **Methods**). Briefly, in CFC and Sham hippocampal recordings, we first detected network bursts of firing across CA1 during NREM sleep. Within these bursts, we calculated the frequency dependent spiking asymmetry between neurons – i.e. whether, statistically, neurons which fire faster during wake also lead spiking during NREM sleep. For each recording, we defined an asymmetry matrix *A*, an *N × N* matrix whose rows and columns were ordered by the relative firing rate of neurons during wake (**Figure 3C**).

We determined the significance, *Z*_*ij*_, of asymmetry for every pair, *A*_*ij*_, and compared the normalized histograms of *Z*_*ij*_ (see representative examples from simulated data, **Figure 3D inset**) when learning was implemented in the model (i.e. an initial engram was present, **Figure 1B**) and when it was not (**Figure 1A**). **Figure 3D** shows the differences between baseline and post-learning distributions of these two cases, showing that distributions after learning are more skewed (with respect to baseline) towards positive Z values than in a non-learning condition (**Figure 3D**). We compared these results to the asymmetry significance distributions observed during NREM for CA1 recordings from experimental CFC and Sham groups, and found similar results (**Figure 3E**; inset shows representative distributions before and after CFC for a representative mouse). These results show that consolidation after learning significantly increases the number of neuron pairs with consistently asymmetric firing patterns during NREM. This indicates the presence of a phase-coding mechanism during NREM sleep, which is sensitive to network changes caused by prior learning.

Finally, to further confirm that the shift in pairwise Z-score distributions is not simply a reflection of frequency differences between the spiking neurons in NREM, we measured global asymmetry across the network during NREM as a function of firing frequency during wake, for every animal separately. To do this we calculated a mean asymmetry score by subtracting the mean asymmetry of the upper triangle and lower triangle of the asymmetry matrix (**Figure 3C**). We then subjected the result to bootstrapping to estimate its significance. We reversed the calculation and repeated the asymmetry calculation for bursts detected during wake, as a function of firing frequency during NREM. These results are depicted in **Figure 3F**, where we plot the significance of the of the NREM asymmetry vs. wake asymmetry. We observe that following CFC (but not Sham conditioning), all animals have higher significances for NREM asymmetry as compared to wake asymmetry.

Based on this relationship, we hypothesized that due to these firing phase differences, STDP during NREM would cause: 1) excitatory connections from high-firing to low-firing neurons to be strengthened, and 2) connections from low-firing to high-firing neurons to be weakened.

### Network oscillations promote temporal coding during NREM sleep

We hypothesize that network oscillations which are naturally augmented during post-learning NREM sleep play a vital role in driving memory consolidation, via STDP. To test this, we next investigated how disruptions of network oscillations affect phase-coding mechanisms. To this end, we prevented inhibitory neurons from firing in the simulated network after learning. Compared to oscillating firing patterns of type 2 networks following learning in the unperturbed condition (**Figure 4A**), this manipulation reduced the oscillatory patterning of excitatory neurons in the network (**Figure 4B**). We compared the effect that this disruption has on post-learning spiking asymmetry (**Figure 4C**). The unperturbed model exhibited stronger positive, significant spiking asymmetry among firing patterns as compared to the one with blocked inhibitory neurons, indicating that oscillations are vital for phase-coding. We also analyzed the spiking asymmetry of CA1 recordings from mice expressing the inhibitory DREADD (Designer Receptor Exclusively Activated by Designer Drugs) hM4Di in parvalbumin-expressing (PV+) interneurons. These mice were treated with either a vehicle (DMSO) or the hM4Di activator clozapine-N-oxide (CNO, to activate hM4Di and suppress PV+ interneuron activity) immediately following CFC. Previous work has shown that CA1 network-wide oscillations and CFM consolidation are disrupted in these mice when CNO is administered after CFC [17, 22]. We found that disruption of PV+ interneuron activity with CNO reduced firing rate-associated spiking asymmetries during post-CFC NREM sleep network bursts relative to DMSO-treated mice, which have normal CA1 oscillations and CFM consolidation (**Figure 4D**). Thus, for both network models in a type 2 regime and the CA1 network during NREM sleep *in vivo*, disruption of oscillations driven by interneurons impairs temporal coding.

**Figure 4:**
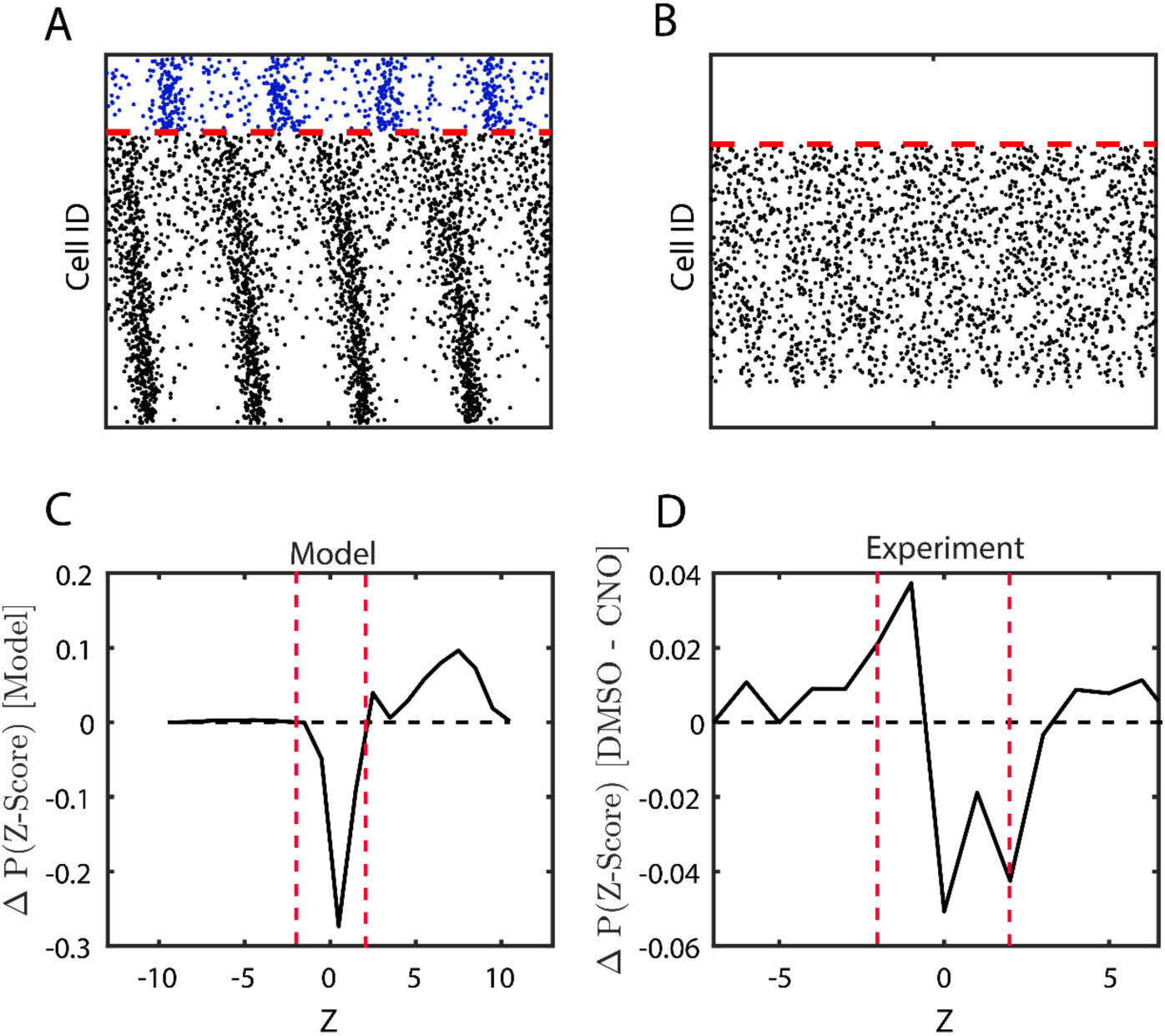
Disruptions to network oscillations lead to diminished temporal relationships between firing neurons. **A, B) SIMULATED DATA:** Simulated raster-plots; **A)** NREM dynamics with learning and before consolidation - full network, and **B)** the same but excitatory cells only with blocked inhibitory cell activity. **C)** Weighted sum of differences in model asymmetry significance distributions after consolidation (unperturbed model in (**A**) minus model without inhibition in (**B**)). **D**) **EXPERIMENTAL DATA:** Difference in experimental asymmetry significance distributions for CA1 recordings for mice expressing the inhibitory DREADD hM4Di in PV+ interneurons, treated with either DMSO (analogous to the full model network) or CNO (reduced network inhibition).

### Frequency-dependent NREM firing asymmetry effects network reorganization through STDP

We next examined whether firing rate reorganization occurs during NREM sleep via STDP in the context of firing asymmetries. In our network model, we compared firing rate reorganization between type 2 excitatory networks with and without firing in inhibitory neurons, when synaptic strength was allowed to evolve over time using an STDP-like plasticity rule. We examined the changes in wake neuronal firing frequencies after an interval in these two scenarios. In the model with normal inhibitory neuron firing, STDP-based synaptic changes in a type 2 (NREM) regime led to a simultaneous increase in the firing rates of principal neurons with the lowest baseline activity and decrease in the firing rates for the most active neurons (**Figure 5A**). We color-coded neurons based on their relative change in frequency across sleep. As shown in **Figure 5A** (inset), neurons which fire faster (vs. slower) at baseline also fire earlier (vs. later) in the oscillation during type 2 dynamics (consistent with **Figure 3**), and experience a decrease (vs. increase) in firing rate due to STDP. Some neurons did not fire during NREM and so did not show a firing frequency change due to STDP (black points in **Figure 5A**). Disruption of network oscillations via inhibitory neuron silencing disrupted the relationship between firing rate changes across the type 2 regime and baseline firing rates for principal neurons (**Figure 5B**). We compared these results with data recorded from the hippocampus of mice with and without DREADD-mediated disruption of PV+ interneuron activity [17, 22]. We measured changes in firing frequency across a six-hour time interval at the start of the rest phase (i.e., starting at lights on), either at baseline (i.e., the day before CFC) or in the hours following CFC. Firing rate changes for each neuron were calculated for mice treated with either DMSO or CNO as a function of their baseline firing rate. The resulting best-fit lines reveal that while CA1 neurons show a relatively low degree of firing rate reorganization across baseline rest, following CFC reorganization is more dramatic. The greatest degree of reorganization is seen after CFC in the control (DMSO) condition, with less-dramatic firing rate changes seen in mice with disrupted CA1 PV+ interneuron activity (CNO) (**Figure 5C**).

**Figure 5:**
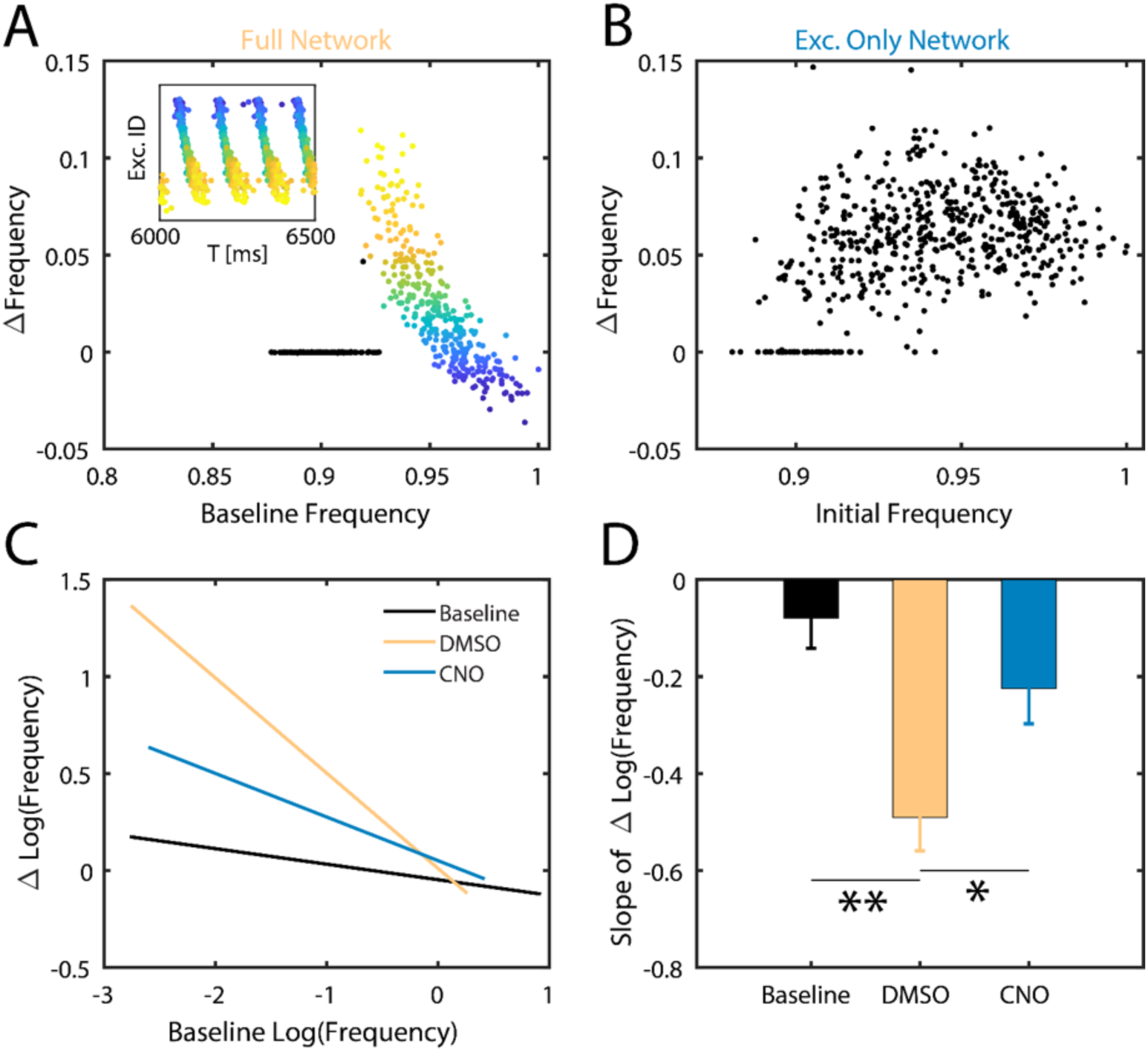
Memory consolidation during sleep differentially affects frequency of firing neurons – model prediction and experiment. **SIMULATED DATA**: **A)** Changes in individual neurons’ firing frequency (normalized to baseline) across post-learning NREM sleep as a function of normalized baseline firing frequency (top). Conditions comparable to **Figure 4A. Inset:** Snapshot of the corresponding raster plot in NREM sleep at the onset of consolidation. Neurons are color-coded based on their change in frequency from baseline and the color is conserved in the raster plot. Black data points (top) are for neurons which did not fire during NREM sleep. Of the neurons that are consistently active, those with initially lower frequency increase their frequency whereas neurons with initially higher frequency decrease their frequency. **B)** Change in firing frequency (normalized to baseline) as a function of normalized baseline firing frequency in the absence of inhibition. Conditions comparable to **Figure 4B.** Unlike the full-network condition (with inhibition), firing frequency changes homogenously across baseline firing rates. **EXPERIMENTAL DATA**: **C)** Best fit lines of the change in log firing rates vs the initial log firing rate, comparing baseline recordings (solid lines; composite n = 11) to post recordings (dashed lines) for the first 6 hours post CFC for mice expressing the inhibitory DREADD hM4Di in PV+ interneurons, treated with either DMSO (Yellow; n = 3; analogous to the full model network) or CNO (Teal; n = 3; analogous to reduced network inhibition), and wild type mice with post-CFC SD (Blue; n = 5). **D)** Slope comparison of change in log firing rates for DMSO, CNO, and SD baseline and post-CFC recordings. Analysis of covariation revealed statistically significant slope differences between DMSO and CNO (*, p = 0.0114), and DMSO and Baseline (**, p < 0.0001).

Comparing the slopes of firing rate vs. firing rate change relationships (**Figure 5D**) for post-CFC recordings, we found significantly weaker reorganization of firing rates in CNO-treated mice relative to DMSO-treated mice. This reorganization of firing rates across the network is an important prediction of the model, as it suggests a possible universal network-level correlate of sleep-based memory consolidation *in vivo*.

## Discussion

Sleep is vital for successful memory consolidation across organisms and different types of memories (e.g., those mediated by network activity in hippocampus vs. sensory cortex [15]). Similarly, recent experimental advances have shown that oscillatory dynamics in neural circuits play a vital role in memory consolidation [22, 32, 33]. Here, we argue that temporal organization of neuronal firing by network oscillations expressed during NREM sleep promotes feed-forward synaptic plasticity (i.e., STDP) from highly active neurons to less active ones. This process in turn promotes recruitment of additional neurons into the engram – the “systems consolidation” underlying long-term memory storage.

Our model demonstrates that the initial encoding of memories in a network (e.g. in the CA1 network during CFC) augments low frequency network oscillations (**Figure 1, 2D**). The augmentation of these oscillations occurs in a NREM sleep-like, low-ACh, type 2 network state. In this state, neurons are less responsive to input (i.e. the input-frequency curve flattens [14]), and they exhibit higher propensity to synchronize to periodic input (i.e., network oscillations). This locking to network oscillations leads to more stable firing relationships between neurons. We observe this increase in stability (i.e. increase in FuNS) both in our hippocampal network model after introduction of a synaptically-encoded memory, and in the mouse hippocampus *in vivo* following single-trial CFC (**Figure 2**).

Our prior work has shown that sleep-associated FuNS changes recorded in the CA1 network following CFC is a salient feature of network-wide dynamics that accompanies successful memory consolidation (**Figures 2-5**) [7, 17, 20, 22]. We find that increased FuNS is associated with stronger low-frequency oscillatory patterning of the network, and predicts which experimental conditions will support, disrupt, or rescue fear memory consolidation (**Figure 2C**). This increased stability in turn mediates mapping between firing rates during wake and relative phase-of-firing during NREM sleep (**Figure 3A, B**) – such that firing of neurons with higher baseline firing frequency leads those with lower baseline firing frequency. We showed that this frequency/timing relationship is also detected in experimental data from CA1 during NREM sleep (**Figure 3C, E, F**). Moreover, when network oscillations are abolished by inhibitory neuron activity (**Figure 4**) the frequency/timing relationship is disrupted, indicating that oscillations do play central role in this process.

Finally, when synaptic strength in the model is allowed to evolve through STDP, we observe reorganization of firing rates across NREM sleep - with sparsely firing neurons increasing their firing rate substantially, and highly active neurons decreasing their firing rate. These results are also observed in CA1 during sleep-dependent CFM consolidation (**Figure 5**). We again observed a disruption of these effects when the oscillatory network activity is reduced via manipulations of interneurons. Similar sleep-associated firing rate changes have been reported in neurons recorded from various neural circuits [6, 7, 34].

Taken together, our results suggest a universal mesoscopic network mechanism underlying what is commonly referred to as systems memory consolidation. They also provide support for the hypothesis that while the brain may use a firing rate-based code during waking experience, firing phase-based information coding in the context of network oscillations in sleep could play an instructive role for memory storage. This mechanism would mitigate the aforementioned limitations of rate-based information coding in the brain.

While the present study is focused on varying parameters of computational models to predict data from the CA1 network during CFM consolidation, we believe that the mechanisms outlined here may be universally true. For example, sleep, and sleep-associated network oscillations, are required for consolidation of experience-dependent sensory plasticity in the visual cortex [18, 33, 35], and disruption of other hippocampal oscillations during sleep disrupts consolidation of other forms of memory [12]. Moreover, similar frequency-dependent changes in neuronal firing rates are also observed across periods of sleep in the visual cortex [7] and frontal cortex [6]. Based on these and other recent data linking network oscillations in sleep to many forms of memory consolidation, this suggests a unifying principle for sleep effects on cognitive function, and one that could reconcile discrepant findings on how sleep affects synaptic strength [15]. It also provides an expanded and possibly alternative explanation of the role of sleep in memory management than what is often proposed [36]. Here we show that NREM sleep facilitates both increases and decreases in neuronal firing rates in the context of network reorganization, while at the same time recruiting heterogeneous neuronal populations during systems memory consolidation.

## Acknowledgements

Support for this research was provided by the NIH (DP2MH104119 and R01EB018297).

## Author Contributions

Experimental design was done by N.O. and S.A. and was performed by N.O., B.C., and J.S. Modeling design was done by Q.S., B.E. and M.Z. and was performed by Q.S. and B.E. Data analysis was performed by Q.S., B.E., B.C., and J.S. The authors Q.S., M.Z., and S.A. wrote the manuscript.

## STAR Methods

### Lead Contact and Materials Availability

Further information and requests for data or code should be directed to, and will be fulfilled by, the Lead Contact, Michal Zochowski.

### Experimental Model and Subject Details

#### Hippocampal recordings, fear conditioning, and sleep deprivation

All procedures were approved by the University of Michigan Animal Care and Use Committee. Male C57BL/6J mice between 2 and 6 months were implanted using methods described previously [35, 37]. Recording implants (described in more detail in [20]) consisted of custom built driveable headstages with two bundles of stereotrodes implanted in bilateral CA1 and three EMG electrodes to monitor nuchal muscle activity. The signals from the stereotrodes were split into local field potential (0.5-200 Hz) and spike data (200 Hz-8 kHz).

Following post-operative recovery, mice either underwent CFC (placement into a novel environmental context, followed 2.5 min later by a 2-s, 0.75 mA foot shock; n = 5 mice), Sham conditioning (placement in a novel context without foot shock; Sham; n = 3 mice), or CFC followed by 6 h of sleep deprivation by gentle handling (a manipulation known to disrupt fear memory consolidation [17, 20, 21, 30]; SD; n = 5 mice). Spike data from individual neurons was discriminated offline using standard methods (consistent waveform shape and amplitude on the two stereotrode wires, relative cluster position of spike waveforms in principle component space, ISI ≥ 1 *ms*) [7, 17, 20, 22, 33]. Only neurons that were stably recorded and reliably discriminated throughout the entire baseline and post-conditioning period were included in subsequent analyses of network dynamics.

24 h following CFC or Sham conditioning, freezing behavior upon return to the conditioning context was measured to evaluate CFM.

#### Pharmacogenetic inhibition of interneurons

2-3-month-old male Pvalb-IRES-CRE mice were bilaterally injected with either the inhibitory receptor hM4Di (rAAV2/Ef1A-DIo-hM4Di-mCherry; UNC Vector Core: Lot #AV4,708) or a control mCherry reporter (raav2/Ef1A-DIo-mCherry; UNC Vector Core: Lot #AV4375FA) (methods further elaborated in [17]). Using the same implant procedures described above, the animals were implanted with stereotrode bundles.

After allowing 4 weeks for viral expression, the animals underwent contextual fear conditioning (as described above). Post-shock, mice were either given an i.p. injection of either 0.3 mg/kg clozapine-N-oxide (CNO) dissolved in DMSO (to activate the DREADD) or DMSO alone (as a control) [22].

### Methods Details

#### Mixed excitatory-inhibitory conductance-based neuronal networks

Conductance-based neuronal networks containing both excitatory and inhibitory neurons were modeled using a modified Hodgkin-Huxley formalism [25, 38]. The time-dependent voltage *V*_*i*_ of a single neuron is given by

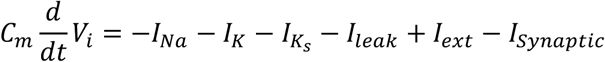

where *C*_*m*_ is the membrane capacitance, *I*_*ext*_ is the fixed external input used to elicit spiking, *I*_*leak*_ = 0.02(*V*_*i*_ + 60) is the leakage current, and *I*_*synaptic*_ = (∑_*j*∈*Excitatory*_ *g*_*E − X*_ *S*_*ij*_)(*V*_*i*_ − *V*_*Excitatory*_) + (∑_*j*∈*Inhibitory*_ *g*_*I − X*_ *S*_*ij*_)(*V*_*i*_ − *V*_*Inhibitory*_) is the total summed synaptic input received by a neuron from its pre-synaptic partners and *g*_*I − X*_ and *g*_*E − X*_ represent the synaptic conductance for connections from inhibitory and excitatory neurons to their post synaptic targets *X* (values provided below). The synaptic reversal potentials are *V*_*Exitatory*_ = 0 *mV* and *V*_*Inhibitory*_ = −75 *mV*. Here, 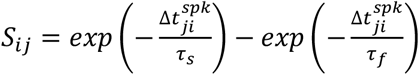 represents the shape of the synaptic current, given the difference in spike timing between the post-synaptic neuron *i* and the recently fired pre-synaptic neuron *j*, 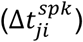, with *τ*_*f*_ = 5 *ms* and *τ*_*s*_ = 250 *ms* or *τ*_*s*_ = 30 *ms* for excitatory synaptic currents and inhibitory synaptic currents, respectively.

The ionic currents are *I*_*Na*_, *I*_*K*_, *and* 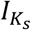, representing sodium (Na), potassium (K), and muscarinic slow potassium (K_s_), respectively. More specifically: 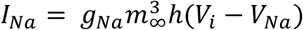, with 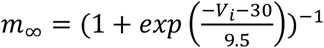 being the activation of the channel and where *h*, the inactivation, is given by the solution to 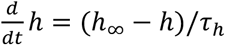, with 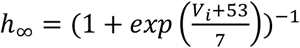 and 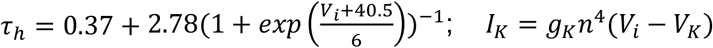 with 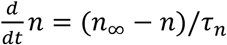 where 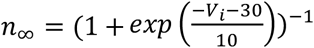 and 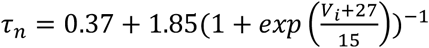; and 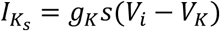 with 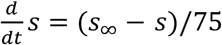 where 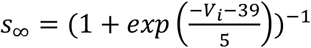 The reversal potentials are *V*_*Na*_ = 55 *mV* and *V*_*K*_ = −90 *mV* and the maximal conductances are 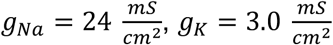.

The slow potassium conductance

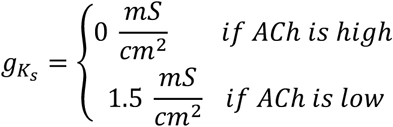

Controls the level of ACh, e.g. during wakefulness (high ACh) or NREM sleep (low ACh). The values thus control the excitability type, where low g_Ks_ (high ACh) yields Type 1 excitability and high gKs (low ACh) yields type 2 excitability. Type 1 excitability is characterized by arbitrarily low firing frequencies, high frequency gain as a function of constant input, and a constant advance in the phase response curve whereas type 2 has a threshold in firing frequency onset, a shallow frequency gain function, and a biphasic phase response curve. See Stiefel, Gutkin, and Sejnowski [25].

Each simulation was completed using the RK4 integration method with a step size of h = 0.05 ms.

#### Network properties

The network used in these studies consists of N=1000 neurons, with *N*_*e*_ = 800 excitatory neurons and *N*_*I*_ = 200 inhibitory neurons. Connections form a random network with different levels of connectivity dependent on the pairwise pre- and post-synaptic neuron identity: Inhibitory neurons project to 50% of the inhibitory neurons and 30% to the excitatory neurons whereas excitatory neurons project to just 6% of both the inhibitory and excitatory neurons, with self-connections being forbidden in all cases. The initial synaptic weights are 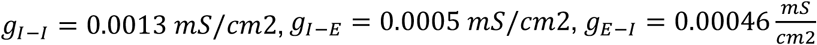, and *g*_*E − E*_ = 0.00003 *mS*/*cm*2 in **Figures 1** and **2**, but with *g*_*E − E*_ = 0.00001 *mS*/*cm*2 in **Figures 3-5**. For initial engram formation, reciprocal connections among a random subset of 250 excitatory neurons increase their *g*_*E − E*_ conductances, constituting the strength of the memory. The effect of the degree of increase in this synaptic strength is investigated in **Figure 2** and elsewhere is kept at 10x.

#### Implementing STDP in the network

Neural correlates of memory are thought to emerge due to the strengthening and weakening of synaptic strengths in an activity-based manner. Here, we use a symmetric learning rule that uniformly increases or decreases synaptic weights based on the time-ordering of pre- and postsynaptic pair firings, only in excitatory-to-excitatory connections. If a presynaptic neuron fires before its postsynaptic partner, the conductance increases by an amount 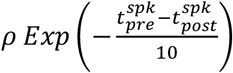. Similarly, a weakening of synaptic strength occurs by an amount 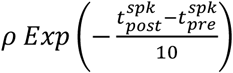 when a postsynaptic neuron fires before its presynaptic partner. In both cases, if the time difference between spike pairs is too great, the change in synaptic strength will approach zero. On the other hand, highly coincident spike pairs will have a maximal change given by the learning rate *ρ* = 10^− 3^. It should be noted that while the synaptic weight is prohibited from becoming negative, there is no upper-bound set on the synaptic strength, though previous work has shown saturation of synaptic weights given sufficient time [14].

#### Analysis of functional network structures through AMD and FuNS

Average Minimal Distance (AMD) [39] was applied to network spiking data to determine functional connectivity. AMD calculates the mean value of the smallest temporal difference between all spikes in one neuron and all spikes in another neuron. Analytical calculations of the expected mean and standard deviation of minimal distance is then used to rapidly determine the significance of pairwise minimal distance [29]. Specifically, the first and second raw moments of minimal distance for each node are calculated: 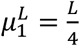 and 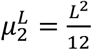, where L is the temporal length of the interspike interval and we have assumed that (looking both forward and backward in time) the maximum temporal distance between spikes is 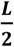. Over the entire recording interval T, the probability of observing an inter-spike interval of length L is simply 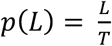. Then, the first and second moments of minimal distance considering the full recording interval are given as 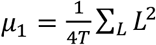 and 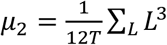. Finally, the calculated statistical moments give rise to the expected mean and standard deviation, *μ* = *μ*_1_and 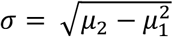, which are used to determine the Z-score significance of pairwise connectivity: 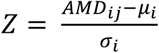. Values of *Z*_*ij*_ ≥ 2 represent significant functional connections between node pairs.

Functional Network Stability (FuNS) tracks global changes in network functional connectivity by quantifying similarities in AMD matrices over a recording interval. The procedure is as follows: first, a recording interval is split into n partitions of equal temporal length. Each partition is subjected to AMD functional connectivity analysis, resulting in n functional connectivity matrices Z. Similarity between time-adjacent functional networks is determined using the normalized dot product after matrix vectorization. FuNS is then determined by taking the mean of these cosine similarities over the recording interval: 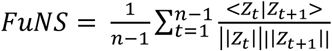. Thus, not only does FuNS yield insight into how functional connectivity changes over time, it can also shed light on how behavior affects the underlying functional network, e.g. by quantifying the difference between FuNS calculated before and after a learning task.

#### Spectral analysis, spike-field coherence, and phase relationships

Histograms of neuronal spiking per unit time were used to calculate the network characteristic frequency, spike-field coherence, and phase relationships of individual neurons to the network signal. First, spike timings were converted into binary spike vectors and then summed to give a network spike vector. Then, the spike vector was convolved with a Gaussian distribution with zero mean and a standard deviation of ∼2 ms, giving a continuous network signal. The spectral power was measured by taking the Fourier transform of the excitatory, non-engram signal (i.e. no neurons with artificially strengthened connections were used; **Figure 1**). Then, the change in spectral power (e.g., in **Figure 2**) was calculated by integrating the frequency-domain signals and taking the relevant percent difference.

Finally, phase relationships of excitatory neuronal spiking compared to inhibitory local field potentials was calculated (please see **Figure 3A**). Peaks in the inhibitory signal acted as the start and end of a given phase and excitatory spike times were used as place-markers of phase between an individual neuron and the signal. The phase of spike occurrence was normalized to give values between 0 and 1.

#### Burst detection

We analyzed bursts of activity for spiking asymmetry between active neurons. First, recordings of a given interval of length L were segmented into smaller windows of length x (25 ms in CFC, 50 ms in CNO/DMSO, and 100ms in model simulations; with times chosen to maximize number of pairwise co-activations occurring) with windows overlapping by 12.5 ms to increase the sampling of the interval L and to reduce effects of windows onset. Then, the total number of **active neurons**, in each window is determined and used to define a burst-detection threshold: a burst occurs if the activity in a window is significantly greater than the mean background activity, averaged over all intervals of a given vigilance state. Specifically, if a window ***w***_***i***_ has a corresponding number of active neurons ***k***_***i***_, then the set of windows representing bursts over all intervals is given as ***B*** = {***w***_***i***_|***k***_***i***_ **≥ *μ***_***k***_ **+ 2*σ***_***k***_}, where ***μ***_***k***_ and ***σ***_***k***_ are the mean and standard deviation across all w.

#### Spiking asymmetry calculation

Next, the pairwise spiking asymmetry A is calculated across all bursts, where A is an *N × N* matrix with entries ***A***_*m,n*_ > 0 if the spikes of neuron m occur before the spikes of neuron n on average, and ***A***_*m,n*_ < 0 in the opposite case. The exact value of an entry ***A***_*m,n*_ is given as the normalized sum of fractional differences between the number of spikes of neuron n occurring after and before each spike of neuron m, across all detected bursts B:

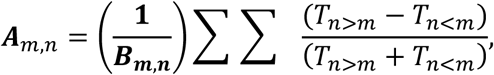

Where *T*_*n>m*_ represents the number of spikes of neuron n occurring after a given spike of neuron m, the inner sum is over the number of spikes of neuron m within a given burst, the outer sum is over all bursts, and the normalization factor is the total number of bursts where neurons n and m are coactive.

#### Relating spiking asymmetry to phase coding of activation frequency

The rows and columns of A are sorted by neuron spiking rate measured within a given vigilance state (i.e. wake or NREM). After sorting, spiking asymmetry of slow firing rate neurons compared to high firing rate neurons will (a) compose the lower triangular matrix of A and (b) will be more positive than the upper triangular matrix of A if faster firing neurons lead slower firing neurons. We thus compared each pairwise entry of *A*_*m,n*_ in the lower triangular matrix with its reciprocal *A*_*n,m*_ in the upper triangular,

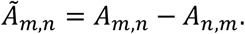

If *Ã*_*m,n*_ > 0, then the faster firing neuron leads the slower firing neuron on average.

We next determined the significance of each *Ã*_*m,n*_ by randomizing the timing of each neuron’s spikes within each burst, 100 times. The value of significance is then given by the Z-score,

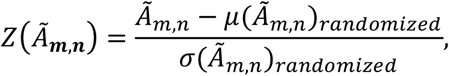

Where *μ*(*Ã*_*m,n*_)_*Randomized*_ and *σ*(*Ã*_*m,n*_)_*Randomized*_ are the mean and standard deviation of the randomized distributions and with *Z*(*Ã*_***m,n***_) ≥ 2 indicating that neuron m leads neuron n in a significant way. In all, we obtain a distribution with 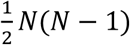 elements, where N is number of detected neurons.

Finally, we compare the changes distributions of *Z*(*Ã*_***m,n***_) for the investigated cases (Figure 3 and Figure 4). A positive difference at *Z*(*Ã*_***m,n***_) ≥ 2 indicates an increase in fast firing neurons leading slow firing neurons within a burst of activity.

Alternatively, instead of calculating the individual pairwise differences in asymmetry, *Ã*_*m,n*_, a global assessment of asymmetry is achieved by first averaging all the individual asymmetry values below the *m=n* diagonal (i.e. ⟨*A*_*m,n*_⟩, where *m* < *n*) and subtracting the average value from above the *m=n* diagonal (i.e. ⟨*A*_*m,n*_⟩, where *m* > *n*). Bootstrapping of spike times within bursts can then be used to determine significance of this difference, with positive values greater than or equal to a value of 2 indicating a significant global increase in fast firing neurons leading slow firing neurons within the network (**Figure 3F**).

